# Augmented expansion of Treg cells from healthy and autoimmune subjects via adult progenitor cell co-culture

**DOI:** 10.1101/2020.12.03.410316

**Authors:** JL Reading, VD Roobrouck, CM Hull, PD Becker, J Beyens, A Valentin-Torres, D Boardman, E Nova Lamperti, S Stubblefield, G Lombardi, R Deans, AE Ting, T Tree

## Abstract

Recent clinical experience has demonstrated that adoptive regulatory T cell therapy is a safe and feasible strategy to suppress immunopathology via induction of host tolerance to allo- and autoantigens. However, clinical trials continue to be compromised due to an inability to manufacture a sufficient Treg cell dose. Multipotent adult progenitor cells (MAPC^Ⓡ^) promote regulatory T cell differentiation *in vitro*, suggesting they may be repurposed to enhance *ex vivo* expansion of Tregs for adoptive cellular therapy. Here, we use a GMP compatible Treg expansion platform to demonstrate that MAPC cell-co-cultured Tregs (MulTreg) exhibit a log-fold increase in yield across two independent cohorts, reducing time to target dose by an average of 30%. Enhanced expansion is linked with a distinct Treg cell-intrinsic transcriptional program, characterized by diminished levels of core exhaustion (*BATF, ID2, PRDM1, LAYN, DUSP1*), and quiescence (*TOB1, TSC22D3*) related genes, coupled to elevated expression of cell-cycle and proliferation loci (*MKI67, CDK1, AURKA, AURKB*). In addition, MulTreg display a unique gut homing (CCR7lo β_7_hi) phenotype and importantly, are more readily expanded from patients with autoimmune disease compared to matched Treg lines, suggesting clinical utility in gut and/or Th1-driven pathology associated with autoimmunity or transplantation. Relative to expanded Tregs, MulTreg retain equivalent and robust purity, FoxP3 TSDR demethylation, nominal effector cytokine production and potent suppression of Th1-driven antigen specific and polyclonal responses *in vitro* and xeno graft vs host disease (xGvHD) *in vivo*. These data support the use of MAPC cell co-culture in adoptive Treg therapy platforms as a means to rescue expansion failure and reduce the time required to manufacture a stable, potently suppressive product.

## Introduction

Regulatory T cells (Tregs) are pivotal regulators of immune responses; maintaining self-tolerance, homeostasis, and controlling excessive immune activation through a spectrum of cell-mediated and soluble mechanisms. The best characterised subset of Tregs are those defined by constitutive expression of CD25 and FOXP3, the master regulator of their suppressive phenotype and function [1]. Tregs play a key role in the prevention of autoimmune diseases, allergies, infection-induced organ pathology, transplant rejection and graft vs host disease (GvHD). Based on encouraging results in pre-clinical models, adoptive transfer of *ex vivo* expanded Tregs is seen as a promising therapeutic strategy to restore immune balance and promote tolerance in individuals undergoing hematopoietic stem cell- and solid organ transplantation or suffering from autoimmune diseases such as Crohn’s disease (CD) and type 1 diabetes (T1D) [2]. Recently, early phase clinical trials have demonstrated that adoptive Treg cell therapy is safe and feasible [3]. However, many clinical trials have been compromised due to manufacturing challenges, primarily in the isolation of pure Tregs and *ex vivo* expansion to produce sufficient cell yields for a clinical dose [3–5].

We recently developed a GMP-compatible platform at King’s College London (KCL) for the isolation and expansion of Tregs for adoptive cell therapy and has been trialed as a method to promote renal allograft tolerance as part of the ONEstudy [6, 7] and the ThRIL study [8, 9]. A key feature of this platform is the inclusion of the mTOR inhibitor Rapamycin, which prevents effector T cell outgrowth [10–12]. In parallel, we have explored the immunomodulatory potential of MAPC cells, an adult, adherent bone-marrow derived stromal cell that is under clinical investigation for numerous indications [13–16] and determined that these cells suppress effector T cell function and promote Treg induction in murine and human models of autoimmunity, transplantation and injury [17–20]. We hypothesized that the immunomodulatory potential of these two clinical grade cell therapies may synergize such that Treg abundance or function could be enhanced in the presence of MAPC cells; thereby establishing superior protocols to advance Treg manufacture for adoptive cell therapy.

## Results

### MAPC cell co-culture results in markedly increased Treg expansion

Based on the potential of clinical grade MAPC cells to promote Treg differentiation, we tested MAPC cell co-culture as a potential strategy to enhance Treg manufacture. To do so, we adopted the GMP compatible Treg process from the ONEstudy and ThRIL as a benchmark protocol [7, 9]. CD4+CD14-CD127loCD25hi live lymphocytes were sorted from freshly isolated PBMC of n=10 healthy volunteers and stimulated 1:1 with anti-CD3/CD28 coated beads and maintained in the presence of 600IU/ml IL-2 and 125ng/ml Rapamycin 10 days, after which cells were replated with fresh anti-CD3/CD28 coated beads (**FigS1A-B)**. Following three, sequential 10-day expansions cells were starved of Rapamycin and IL-2 for 48h, then harvested for endpoint analysis (cells expanded under these conditions hereafter referred to simply as ‘Tregs’). In parallel, we replicated these conditions with the addition of single donor, allogeneic MAPC cells at a ratio of 1:10 MAPC:Treg at day zero of each round. Treg cells generated under these conditions are hereafter referred to as ‘MulTreg’(**Fig.1A**). At day 10 the number of MulTreg cells was 3.5 times greater than paired Tregs (21.4 vs 6.1 fold expansion vs *ex-vivo*). This difference in yield increased to 9.6-fold by day 20 (440 vs 45.9-fold expansion) and further to 15.9-fold upon completion of the expansion at day 30 (11,178 vs 699.6-fold). The greater abundance of MulTreg cells was significant at all timepoints measured (q=3.910^−3^, Multiple Wilcoxon signed rank test), **Fig.1B**. The yield at day 30 was greater (9/10 donors) or equivalent (1 donor) in MulTreg relative to Treg across the cohort (range 0.9-733-fold greater expansion in MulTreg vs Treg), **Fig.1C**.

**Figure 1.**
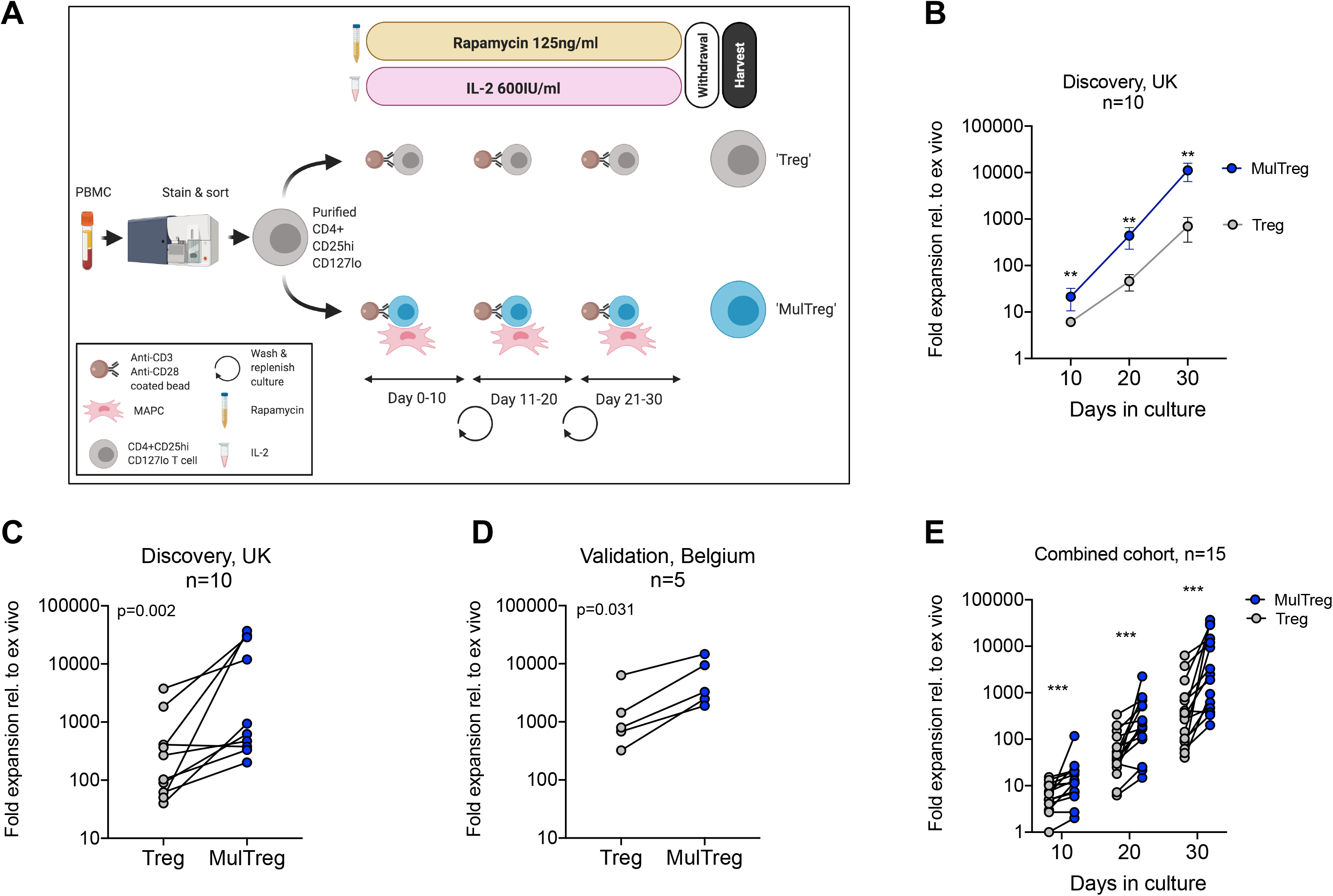
Treg and MulTreg isolation and expansion. (A) Schematic of the expansion protocol for Treg and MulTreg. (B)The fold expansion of Treg and MulTreg lines vs *ex vivo* (fresh Tregs plated post sort) was calculated for each donor at each time point. Datapoints represent the mean +/−SEM for 10 donors. (C)Fold expansion (from *ex vivo*) at d30 for the UK (KCL) cohort (Treg and MulTreg lines grown from 10 individuals). (D)Average fold expansion (from *ex vivo*) at d30 for the validation (Belgian) cohort (Treg and MulTreg lines grown from 5 individuals). (E)Fold expansion in MulTreg versus Treg expansion was calculated for each donor at each time point in the combined cohort: Treg and MulTreg lines grown from 15 individuals. All p and q values from single or multiple (corrected) matched-pairs Wilcoxon-signed rank test. **q<0.01, ***q<0.0005

To validate these results, a separate research team at a commercial cell therapy facility in Leuven, (Belgium, EU) performed expansions in a second, independent validation cohort of 5 donors. Data produced at the Belgian research site confirmed that of the KCL team, demonstrating increased yield in 5/5 donors in MulTreg compared to Treg (range 2.3-7.7-fold, p=0.0313), **Fig1D**. This effect was found to be highly reproducible across 10/10 expansions run from a single donor to assess consistency (p=1×10^−3^ average 8.2 fold, range 1.7 to 20-fold), **FigS1C,** or when using three independent MAPC cell donors (**FigS1D**). Combined analyses from both cohorts showed a significant, consistent and marked increase in MulTreg yield relative to Treg at all timepoints (q=1.2×10^−4^), resulting in an average 9.4-fold increase in quantity by day 30, **Fig1E**. Indeed, MulTreg yields at day 20 were not statistically different from Treg yields at day 30 (p=0.094) suggesting equivalent dose could be reached 10 days earlier via the MulTreg platform. These data indicate that MAPC cell co-culture consistently results in a significantly more rapid expansion and greater yield of Treg cells during *in vitro* GMP-compatible platforms that emulate clinical manufacture protocols.

### MulTreg exhibit stable Treg lineage identity and a CCR7lo Integrin β_7_ hi phenotype

Cells harvested at day 30 were analysed by flow cytometry. Treg and MulTreg cells showed equivalent purity of CD3+CD4+ T cells (mean Treg 96.9+/−3.1% vs MulTreg 97.25+/−2%, p=0.25, **Fig.2A-B**) that were >98% FoxP3 positive (mean Treg 99.35%+/−0.4, MulTreg 99.55%+/−0.2, p=0.44, **Fig.2C**) and expressed equivalent levels of FoxP3 MFI(mean Treg MFI 3366+/−1654 vs MulTreg 3631+/−1157, p=0.56,) which were higher than Teff, as anticipated (MFI 230.5+/−87), **Fig.2D**. These data were independently validated by the Leuven team in the validation cohort (Supplementary **Fig.S2A**, n=5). FoxP3 TSDR methylation analysis showed that, in contrast to Teff (mean methylation 89.9%+/−0.64) expanded Treg (27.4%+/−13.2) and MulTreg (29.25+/−13.6) cells were stably committed to the Treg lineage (**Fig2E**). Correspondingly, the frequency of effector (IFN-gamma, IL-17A, TNF-alpha, granzyme B) and regulatory (IL-10) cytokines produced upon PMA/Io restimulation were low, and comparable between Treg and MulTreg (p=ns for all, **Fig2.F**, Supplementary **Figure S2B**). These data demonstrate that MAPC cell-co-culture generates enhanced yields of highly pure and stably committed regulatory T cells.

**Figure 2.**
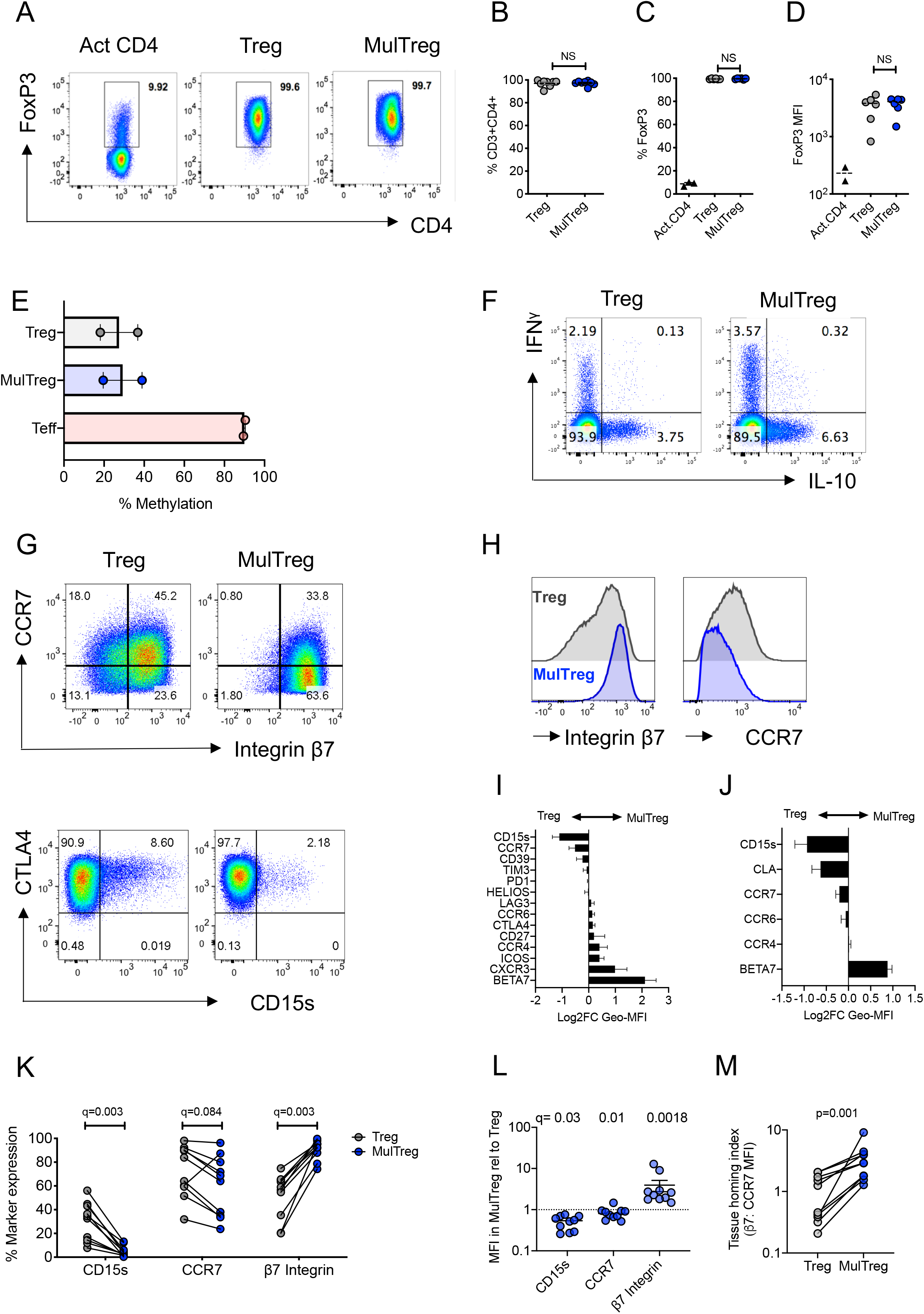
MulTreg are stably committed FoxP3hi CD4+ Treg cells exhibiting a distinct tissue homing phenotype. (A) Example of FoxP3 staining in activated CD4 T cells and Treg or MulTreg lines. (B) Frequency of CD4+ T cells in live gated events, n=8 pairs. (C) Frequency of CD4+ T cells expressing FoxP3 in activated bulk CD4 T cells (n=2) Treg and MulTreg cell lines, n=6 pairs. (D)Median fluorescence intensity of FoxP3 staining in activated CD4 T cells (n=2), Treg and MulTreg cell lines, n=6. (E) Mean percentage of methylation on 9 CpG islands of the *FOXP3* Treg-specific demethylated region (TSDR), n=2. (F) Representative flow cytometry plots of expanded lines following restimulation for 5h with PMA/Io and intracellular staining with the cytokines indicated, gated on live CD4 T cells. (G-H) Representative flow cytometry plots (G) or histograms (H) of expanded lines showing expression of the markers indicated. (I-J) Relative MFI of markers indicated amongst Treg and MulTreg lines (expressed as log2FC in favour of MulTreg) from (I) n=5 donors in the KCL (Discovery) cohort and (J) n=5 donors in the Belgian (Validation) cohort. (K-M) Combined analysis of KCL and Belgian cohorts (n=10) showing frequency (K) or relative MFI (L) of marker expression indicated and the ratio of β_7_ integrin to CCR7 MFI, defined as the tissue homing index (M). Data points represent individual donors, Error bars = SEM of 5 or 10 (L) donors. Stats derived from single or multiple Wilcoxon matched-pairs signed rank test.

The chemokine receptor profile of Tregs directs homing to inflamed tissues, whilst inhibitory receptor and differentiation markers co-define regulatory potential. We therefore assessed Treg differentiation, functional and homing markers in the KCL cohort. Of all markers tested the lymph node homing receptor CCR7 and CD15s (sialyl lewis x, expressed by Tregs in sarcoidosis) were lower in MulTreg, whilst gut homing integrin beta 7 (β_7_) showed a trend for increased expression in MulTreg according to both MFI and frequency(**Fig2G-I**, Supplementary **Fig.2C**). Inhibitory receptors (e.g. CTLA-4, ICOS, PD-1) differentiation markers (e.g. Helios, CD27) and other chemokine receptors (e.g. CCR4) showed no difference in expression (supplementary **Fig.2C**). The Belgian research group independently validated and extended phenotypic analysis in the second cohort, illustrating the same trends for CD15s, CCR7 (both lower in MulTreg) and integrin β_7_ (higher in MulTreg), **Fig.2J**, Supplementary **Fig.S2D**. In addition, the skin homing marker CLA examined by the Belgian team was lower in MulTreg cells (Supplementary **Fig.S2D**). Combined analysis from the two cohorts (n=15) confirmed that the frequency (**Fig.2K**) and/ or MFI (**Fig.2L**) of CD15s and CCR7 was significantly lower-whilst the gut homing integrin β_7_ was significantly and markedly higher in MulTreg cells. Finally, we derived a Treg tissue homing index from the ratio of MFI of β_7_:CCR7 which was significantly higher in MulTreg compared to Treg (p=0.001, **Fig.2M**). These results demonstrate that MAPC cell co-culture generates pure, stably committed Tregs which maintain the expression profile of major inhibitory receptors and exhibit a distinct CD15s^−^ CCR7^lo^β_7_^hi^ phenotype.

### MAPC cell co-culture leads to transcriptional rewiring of exhaustion vs replication related circuitry in expanded Tregs

We next examined the transcriptional profile of the two cell products in an effort to understand the molecular mechanisms underpinning increased expansion and altered phenotype in MulTreg. RNAseq analysis revealed distinct transcriptional programs in Treg vs MulTreg pairs (n=4) evident by clustering on PCA (**Fig.3A**) and heatmap analysis (**Fig3.B)**, which represented a total of 222 significantly differentially expressed genes (DEG). Consistent with growth analysis, MulTreg cells expressed significantly higher levels of genes driving proliferation (*MKI67*, *AURKA*, *AURKB*, *CDK1*), or marking activated cells (*CD38*), whilst negative regulators of cell cycle/activation (*TOB1*, *TSC22*, *KLF2*, *SAMSN1*, *DUSP1*) and genes involved in T cell exhaustion (*LAYN*, *ID2*, *IL27RA*, *NFIL3*) were diminished **Fig.3B-D**. Tregs also displayed higher levels of progenitor and lymph node-homing genes (*CCR7*, *SELL*, *LEF1*), consistent with phenotyping analysis **Fig.3B-D**. Interestingly, *SELL* and *RGS1* (both up-regulated in Treg) are preferentially expressed in sorted β_7_ integrin negative memory CD4 T cells, whilst *ITGA4* (up-regulated in MulTreg) marks β_7_ integrin positive cells, suggesting the expression of β_7_ integrin by flow cytometry reflects a shift in gene expression characteristic of β_7_ integrin-expressing CD4 T cells [21] **Fig.3B-D**. We next assessed whether Treg or MulTreg were enriched for signatures of CD4 T cell subsets isolated from lymphoid or non-lymphoid tissues in inflammatory or homeostatic conditions by gene set enrichment analysis (GESA). Compared to MulTreg, Tregs were significantly enriched for murine signatures of resting/naïve-like regulatory T cells [22] lymphoid resident memory CD4+ T cells [23] and TCF7+ memory progenitor cells [24], reinforcing that MulTreg cells were comparatively skewed towards an activated, more differentiated/tissue homing state (**Fig.3E, Fig.S3A**). To further explore the observation that MulTreg cells were characterized by a less exhausted molecular profile, we examined a shortlist of transcription factors recently shown to define chronically stimulated Tregs that acquired features of T cell exhaustion accompanied by loss of *in vivo* suppressive function [25]. MulTreg cells showed a trend towards lower levels of *TOX*, *BATF*, *ID3*, *PDRM1* (Blimp-1), and *NFKB2* as well as the Th1 and memory associated transcription factors *TBX21* (T-Bet) and *BCL6*, whilst decreased levels of *ID2* and *NFIL3* were found in unsupervised analyses (**Fig.3F**). Notably, three of the four core transcription factors shown to most specifically identify exhausted Treg (vs control Tregs) in the study were each significantly lower in MulTreg compared to Treg (*ID2*, *BATF*, *PDRM1*; p<0.05, ID3 p=ns). These data suggest that the enhanced expansion potential and altered phenotype of MulTreg cells may arise from a selective pattern of transcriptional reprogramming characterized by a shift in the balance of master regulators of T cell quiescence (e.g. *TOB1*, *TSC22D3*), progenitor identity (e.g. *LEF1*), homing (*CCR7*, *SELL*, *ITGA4*, *RGS1*), exhaustion (e.g. *ID2*, *BATF*, *PDRM1*) and replication (e.g. *MKI67*, *AURKA*, *AURKB*, *CDK1*).

**Figure 3.**
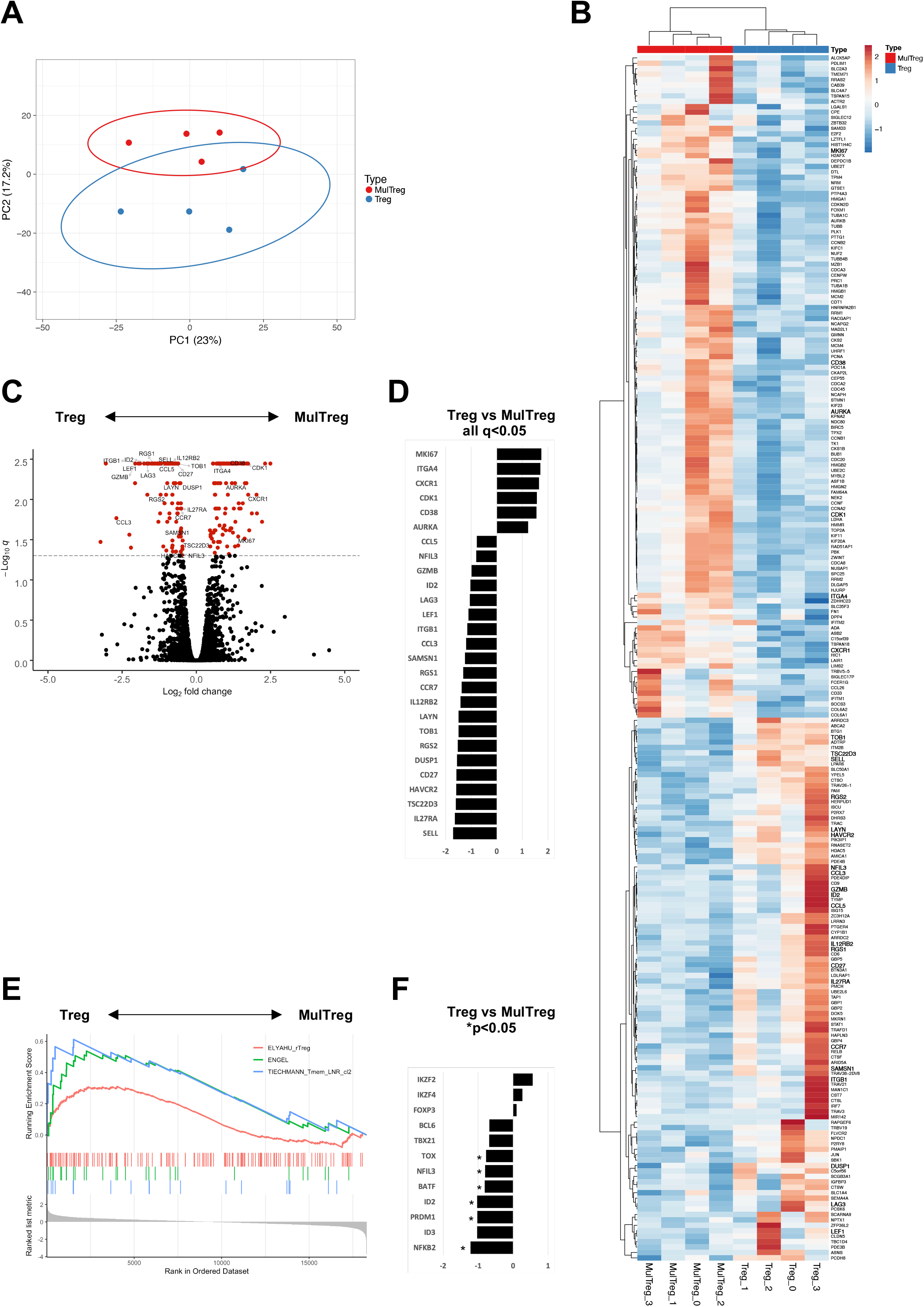
RNAseq analysis of Treg and MulTreg. Pairs of Treg and MulTreg expansions from n=4 healthy donors were analysed by bulk RNAseq. (A) Principal component analysis (B) Heatmap and (C) volcano plot of 222 differentially expressed genes (DEGs) (q<0.05), genes of interest highlighted. (D) Bar plot showing log2-FC of genes of interest amongst DEGs. (E) Enrichment plot from gene set enrichment analysis (GSEA) showing specific enrichment of TCF7+ progenitor memory CD4+ T cells (ENGEL), resting/naive-like Tregs (ELYAHU) and lymph node resident (TIECHMANN) signatures amongst Treg (vs MulTreg) lines. (F) Targeted analysis of Treg exhaustion-related transcription factors within Treg and MulTreg pairs expressed as log2-FC in favour of MulTreg (*p<0.05).

### MulTreg cells suppress human antigen-specific Th1- and polyclonal responses *in vitro*

Polyclonal *in vitro* suppression assays remain the gold standard release criteria for Treg cell therapy products, whilst the majority of immunopathology associated with autoimmunity and graft rejection is elicited by Th1/Th17 antigen-specific responses. We therefore evaluated the *in vitro* suppressive potential of Treg and MulTreg lines using a combination of polyclonal (anti-CD3/CD28 coated beads) and antigen specific, Th1 driven recall (Flu hemagglutinin vaccine ‘Flu-HA’) models. Both lines significantly suppressed T cell proliferation in polyclonally activated 3^rd^ party PBMC (n=5), **Fig.4A-C**. During the polyclonal response, MulTreg co-culture lead to significantly lower IFNg production and MulTreg- but not Treg co-culture lead to significant impairment of TNFa and IL-17A accumulation, whilst both lines inhibited IL-13 secretion (**Fig.4C**). The expansion of CD4 and CD8 T cells to Flu-HA was also attenuated by both Tregs and MulTregs, with a trend for MulTreg to exhibit greater levels of suppression at 2 out of the 3 Teff: Treg ratios in CD4 and CD8 (**Fig.4D-E**). Whilst both cell products showed a clear trend towards inhibition of IFNg and TNFa in Flu-HA responses this was only significant in MulTreg co-cultures, however, no difference was seen in inhibition of IL-13 or modulation of the low levels of IL-17A generated (**Fig4.F**). In addition, MulTreg and Treg cells both suppressed responses to autologous, purified Teff CD4+ T cells (**Fig.S4**). These data highlight that augmented expansion and phenotypic/transcriptomic differences in MulTreg are accompanied by potent *in vitro* suppressive potential, that is equivalent to or greater than that observed with paired Tregs.

**Figure 4.**
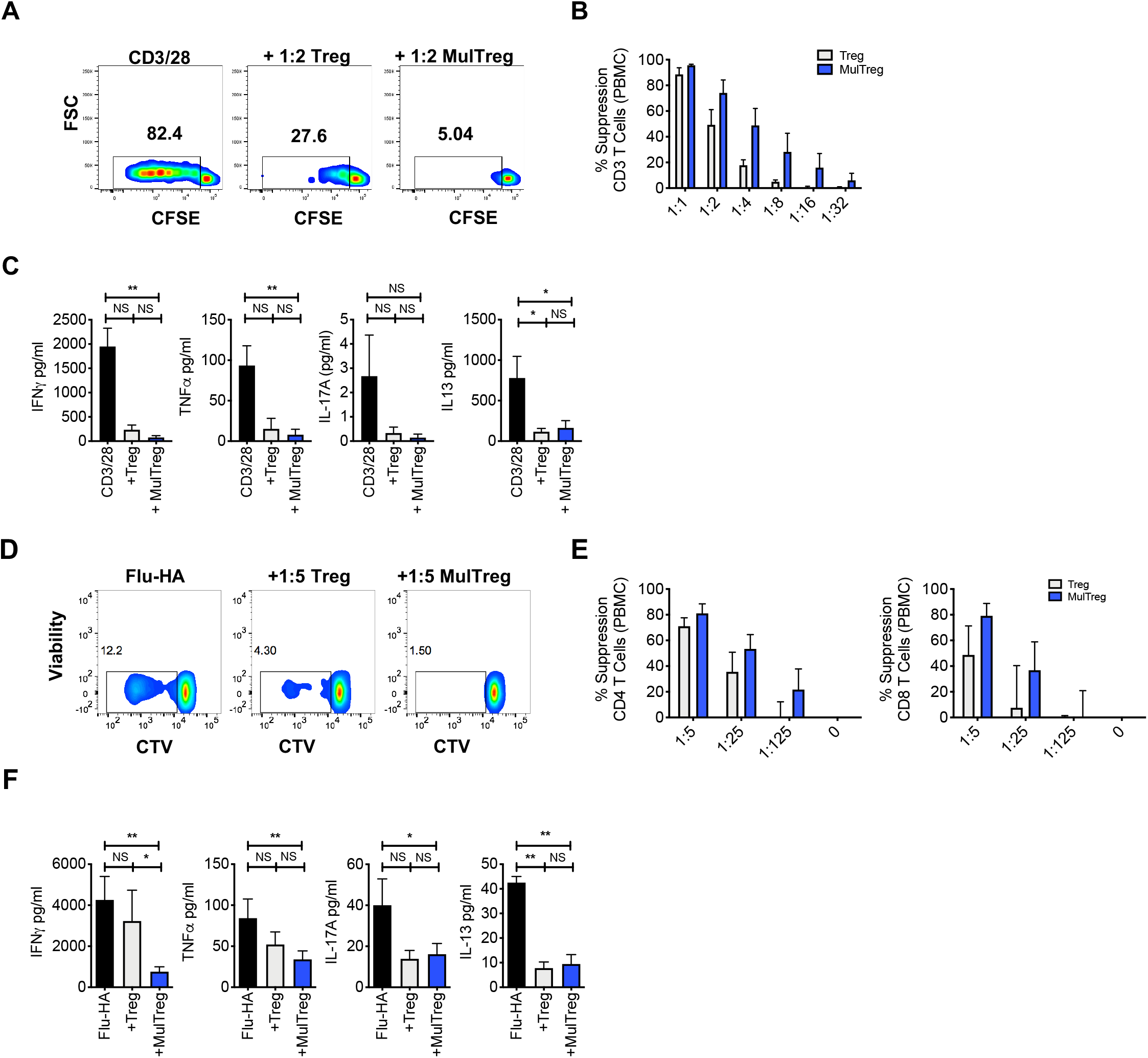
MAPC expanded Tregs exhibit superior suppression of antigen specific responses. (A-C) Suppression assay using 3rd party PBMC stimulated with CD3/CD28 microbeads for 6 days in the presence or absence of Treg or MulTreg lines. (A). Flow cytometry plots showing proliferation in responder T cells from PBMC stimulated with CD3/CD28 microbeads (1:10 bead to cell) in the presence or absence of autologous, expanded Treg or MulTreg lines at a ratio of 1:2 Treg:PBMC. (B) Bar graph displaying % suppression of responder T cell proliferation at different ratios of Treg or MulTreg to PBMC. (C) Bar graph displaying cytokine levels in tissue culture supernatant from suppression assays in panel B (ratio 1:2 Treg:PBMC). n=5 (D-F) Suppression assay using autologous PBMC stimulated with Flu-HA for 6 days in the presence or absence of Treg or MulTreg lines. (D) Flow cytometry plots showing proliferation in responder CD4 T cells from PBMC stimulated with Flu-HA in the presence or absence of autologous, expanded Treg or MulTreg lines at a ratio of 1:5 Treg:PBMC. (E) Bar graph displaying % suppression of responder CD4+ T cell proliferation at different ratios of Treg:PBMC, CD4 (left) and CD8 (right). F) Bar graph displaying cytokine levels in tissue culture supernatant from suppression assays in panel E (ratio of 1:5). Error bars represent the SEM of 5 donors. Stats from Wilcoxon matched-pairs analysis (B,E) or Friedman test (C,F). *p<0.05, **p<0.01

**Figure 5:**
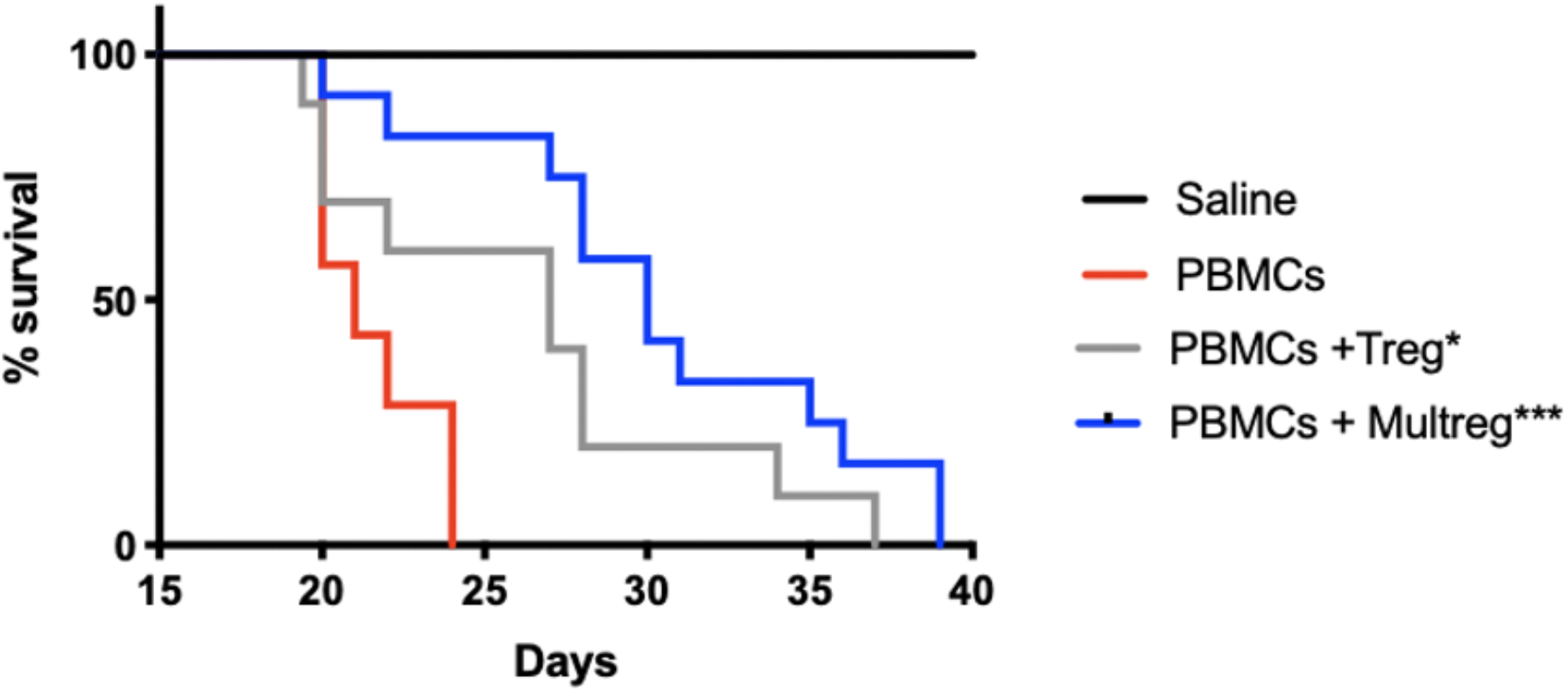
MulTreg effectively control the immune response in a humanised mouse model of xeno-GvHD. NSG mice were inoculated with 10^7^ PBMCs +/− 10^7^ Tregs or MulTregs or PBS as a control (PBMC alone n=7, PBMC and Treg n=10, PBMC and MulTreg n= 12 mice, PBMC and PBS n= 5). Survival of mice administered with MulTreg and Treg in addition to PBMC was significantly longer than those given PBMC alone. Log-rank Mantel-Cox Test (*p<0.05, ***p<0.001).

### MulTreg suppress pathogenic human effector responses during xGVHD *in vivo*

To benchmark the *in vivo* regulatory capacity of MulTreg we assessed their ability to suppress disease in a model of human into mouse xenogeneic GvHD. In this model GvHD is driven by expansion of human T cells, which rapidly adopt an effector memory phenotype, a process that is dependent on xeno-reactivity with foreign MHC class-I and class-II molecules and resembles alloreactivity in the human transplant setting [26]. The engrafted cells can be regulated by therapeutic manipulation making this a suitable model to test the suppressive capacity of Treg cell lines. In the absence of Treg administration disease presented rapidly with no animals surviving beyond 24 days. Administration of Treg or MulTreg significantly slowed the progression of GvHD when compared to mice given PBMCs alone (median survival 27d Treg vs 21d PBMC; p=0.032 and 30d MulTreg vs 21d PBMC; p=0.003) with a trend observed towards a superior survival with MulTreg compared to Treg (median survival 30d MulTreg vs 27d Treg; p=0.1).

### MAPC cell co-culture facilitates superior expansion of suppressive, stable, β_7_hi Treg cells from patients with autoimmune disease

CD4+CD25hiCD127lo cells were sorted from PBMCs of patients with Crohn’s disease (n=4) and T1D (n=2) and expanded using the MulTreg protocol (patient demographic in **Supplementary Table 1**). MulTreg were more readily expanded from autoimmune patients compared to Treg (mean 2159 vs 8239 fold relative to *ex vivo*, p=0.0156), **Fig.6A,** retained equivalent purity and expressed the characteristic CD15slo CLAlo β_7_ hi phenotype (**Fig.6B**). As seen in healthy donors, Treg and MulTreg cells from CD patients harboured a demethylated TSDR (**Fig.6C**) and exerted suppression of polyclonal responses in PBMC according to proliferation (**Fig.6D**) and effector cytokine suppression (**Fig.6E**). These data suggest that MAPC co-culture can be used to rapidly expand a greater yield of suppressive, stably committed Treg cells from populations of patients with prototypic autoimmune disorders, from whom Treg expansion remains a challenge.

**Figure 6:**
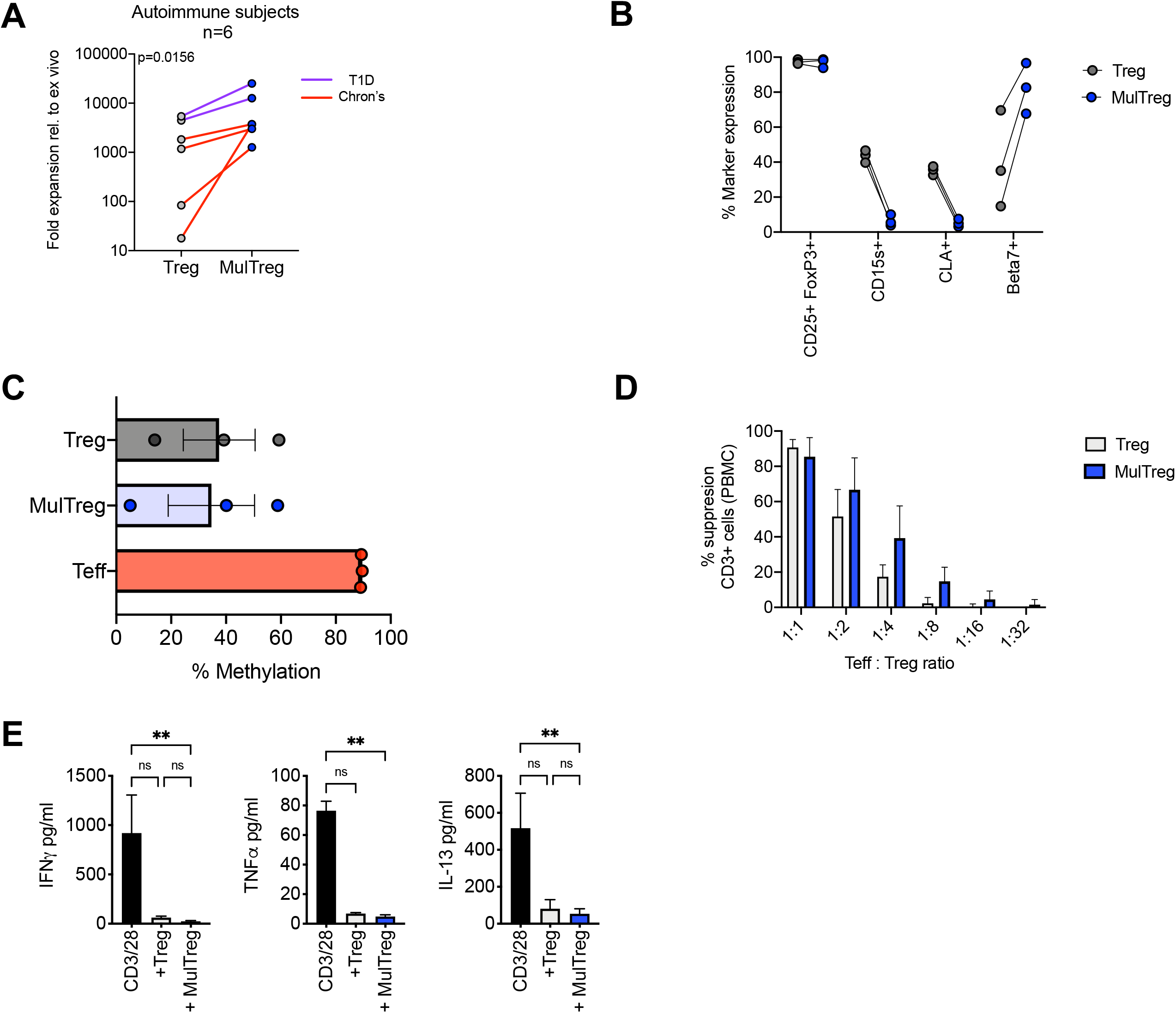
Characteristics of MulTregs from patients with Crohn’s Disease. A) Average fold expansion (from *ex vivo*) for Treg and MulTreg lines grown from 4 individuals with Crohn’s disease and 2 individuals with type 1 diabetes (T1D). Wilcoxon matched-pairs signed rank test (n=6) B) Paired flow cytometric analysis of marker frequencies in Treg and MulTreg cells (n=3) C) Average percentage of methylation on 9 CpG islands of the FOXP3 Treg-specific demethylated region (TSDR) (n=3). Friedman test (ns) (D-E) Suppression assay using 3rd party PBMC stimulated with CD3/CD28 microbeads (1:10 bead to cell) for 6 days in the presence or absence of Treg or MulTreg lines before analysis at d6. (D) Bar graph displaying % suppression of responder CD3+ T cell proliferation at different ratios of Treg:PBMC (n=4). (E) Bar graph displaying cytokine levels in tissue culture supernatant from suppression assays in panel D (ratio 1:2) (n=3), Friedman test. Data in A-D analyzed via Wilcoxon matched pairs analysis ns unless stated.

## Discussion

The past decade has seen intense interest in evaluating the clinical utility of Tregs for a variety of indications including hematopoietic stem cell transplantation, autoimmune diseases and solid organ transplantation [3, 27]. While the results from these initial clinical trials have demonstrated the potential therapeutic value of Tregs, isolation and expansion of Tregs is a major challenge due to reduced Treg frequencies and/or functionally defective Tregs reported in patients with autoimmune disorders [28–34]. In order to overcome this deficiency, a variety of ex vivo strategies have been developed to select, isolate and expand Tregs [4]. Although progress has been made there is still a significant need to develop more robust methods to manufacture clinical grade Tregs.

Numerous *in vitro* and *in vivo* studies have demonstrated the MAPC cells modulate uncontrolled immune responses and increase the proliferation of regulatory T cells [17, 18, 35, 36]. Furthermore, a transient increase in Tregs was also observed in a clinical study evaluating the administration of MultiStem®, a clinical grade product of MAPC cells, in patients receiving a liver transplant [16].

When MAPC cells were introduced into a GMP compatible Treg expansion platform, the resulting MulTregs exhibited greater expansion than paired Tregs and maintained a similar stable and suppressive Treg identity as demonstrated by FoxP3 expression, TSDR methylation status, and the ability to reduce both antigen specific autologous (Flu-HA;**Fig.4B**) and polyclonal T cell proliferation (CD3/28) of autologous (**Fig.S4**) Teff or 3rd-party PBMC **(Fig.4A**). The functional activity of MulTreg was further evaluated in a mouse model of acute xGVHD where it significantly delayed the xGVHD and extended survival of the animals compared to Tregs. Though differences between the two lines were non-significant in proliferation and xGVHD assays, we observed a trend towards greater potency of MulTreg both *in vitro* and *in vivo*.

An emerging issue in T cell therapy is exhaustion due to prolonged TCR stimulation during the manufacturing process and/or *in vivo* chronic antigen exposure [25]. T cell exhaustion is associated with loss of clinical activity in CAR-T cells and tumor-infiltrating lymphocytes (TILs) and a lack of *in vivo* function in Tregs, highlighting a need to generate products less prone to acquiring this hypofunctional state after transfer [37, 38]. Multiple genes coordinate T cell exhaustion [39]. In particular, master transcription factors *TOX*, *PRDM1*, *ID2*, *BATF* play an essential role [40, 41]. A significantly decreased level of expression in these loci may therefore be considered a desirable feature for T cell therapies and could prevent long term loss of function. Compared to Tregs, MulTregs showed significant decrease in levels of three of these four core transcription factors. These data suggest that a faster and more reliable ability to reach the desired Treg dose could potentially be coupled with more durable activity using the MulTreg protocol.

For Tregs therapy to be optimal, infused cells need to migrate to inflammatory sites where they can be activated in the target tissue [42]. β_7_ integrins have been implicated in intestinal T cell homing and retention [43]. Thus, together with loss of *CCR7*, *SELL* (lymph-node homing) and CLA (skin homing), the increased expression of β_7_ integrin on MulTreg suggests a globally altered trafficking profile that may result in preferential gut homing, suggesting clinical utility for the treatment of an autoimmune disorder of the gut. Therefore, we decided to evaluate the Tregs and MulTregs isolated from patients with CD; a debilitating autoimmune disorder of the gut that results in chronic inflammation. Treg treatment is being clinically evaluated in CD [42], however, isolation of Tregs from CD patients results in lower Treg numbers compared to healthy donors, which further supports the need to develop more robust expansion protocols for Tregs isolated from this patient population [42, 44]. Similar to the results with healthy Treg donors, MulTreg isolated and expanded from CD patients had greater expansion potential and higher levels of β_7_ integrin expression but similar FoxP3 methylation expression, and suppression of polyclonal stimulated T cell proliferation compared to Tregs.

Type 1 diabetes is another prototypic autoimmune disorder mediated by Th1 CD4 T cells and CTLs [45] where cell therapy approaches including MAPC cells [17, 18] and Tregs [46–48] show promise. The ability for MulTreg to suppress Th1-driven autologous, antigen-specific recall responses elicited by Flu-HA suggest that these cells can modulate key events in T1D immunopathology [49]. Furthermore, similar to patients with CD, MulTreg cells appear more readily expandable from patients with T1D. Recent reports show that infused Treg cells decline from the circulation of patients with T1D at a highly variable rate, which has been suggested to result from trafficking, turnover or exhaustion [25, 47], implying that reduced exhaustion, well-defined homing profiles or increased cell banks for repeat dosing could be beneficial properties of an adoptive Treg therapy in T1D; highlighting a potential utility of MulTreg in T1D patients.

CAR technology is currently being explored to enhance Treg specificity and functionality [5]. Current CAR-Treg manufacturing approaches implement the initial steps of conventional polyclonal Treg expansion, rendering CAR-Treg manufacturing susceptible to similar manufacturing challenges [50]. Given the promising results in polyclonal Treg expansion in the presence of MAPC cells, we believe that this approach could also be beneficial for CAR-Treg and antigen specific Treg manufacturing.

In summary, the persisting challenges in adoptive T cell and Treg cell therapy include expansion failure from target patient populations, exhaustion, lack of purity and heterogeneity in cell product phenotype. MulTreg cells can potentially be developed as a therapeutic for autoimmune disorders including Crohn’s Disease, offering a means to rapidly expand a stable, less exhausted cell product with a defined, disease-relevant homing phenotype. Importantly this can be achieved via addition of a clinical-grade off the shelf cell product to an existing GMP compatible process which has already demonstrated safety and shows preliminary efficacy.

## Methods

### MAPC cells

MAPC cells used throughout the majority of this study were manufactured by Athersys (Cleveland, OH) using femoral bone marrow aspirates from fully consented donors and processed according to previously described methods [Boozer et al, Journal of Stem Cells, 2009]. The cells were subjected to several quality control tests to guarantee the quality of the expanded cell product (post-thaw viability, flow cytometric analysis of positive/negative surface markers, cytokine secretion for their angiogenic capacity and a T cell proliferation assay to evaluate their immunosuppressive function). MAPC cells were isolated from the bone marrow of a healthy volunteer after obtaining informed consent in accordance with the guidelines of an Institutional Review Board.

### Primary cell culture

PBMC were isolated from fresh blood of consented healthy volunteers by density gradient centrifugation (Lymphoprep, Axis Shield, Oslo, Norway). Stored PBMC samples from individuals with recent onset T1D and HD were included in this study and was approved by the UK National Research Ethics Service (REC# 08/ H0805/14). Written informed consent was obtained from all participants prior to inclusion in the study. Frozen PBMCs from Crohn’s disease patients were obtained from Precision for Medicine or Stem Cell Technologies (Suppl Table 1). Effector (hereafter Teff, CD25^lo^) and regulatory (CD127^lo^ CD25^hi^) T cells were 2-way sorted from PBMC pre-gated on CD4^+^ CD14^−^ viable lymphocytes using a FACS Aria (BD) and the antibodies listed in supplementary material procured from Biolegend or BD. To establish Treg lines, CD127^lo^ CD25^hi^ CD4^+^ lymphocytes were plated at 5×10^4^/well of round bottom 96-well plates (Corning) in 200μl 0.2μM filtered (Sartorius, Terumo) complete media consisting of X-vivo 15 media (Lonza) containing 100μg/ml Penicillin-streptomycin, 100μg/ml amphotericin B (both from Sigma Aldrich), 125ng/ml Rapamycin (Rapamune, Pfizer), 600 IU/ml IL-2 (Proleukin, Chiron), and 5% heat inactivated human AB serum (Sigma Aldrich) and incubated at 37 °C, 5% CO2. Cells were stimulated 2:1 beads to cell with CD3/28 (Dynabeads, Life Technologies) for 72h then harvested, pooled, and transferred to flat bottom 96 well or 48 well plates at a density of 1×10^6^/ml. Cells were maintained by feeding with complete media 2-3 times during days 3-10 and transferred to T75 or T125 when cell numbers exceeded 3×10^7^. This 10-day expansion was repeated twice for a total of 30 days in culture. After 30 days residual beads were removed via magnet (Dynal, Invitrogen), cells were then washed once in X-vivo 15 media and replated at 2×10^6^/ml in X-vivo 15 media containing 2.5% heat inactivated human AB serum for 48h (withdrawal). Autologous MulTreg lines were generated in parallel from the same suspension of sorted Tregs under identical conditions with the exception of adding 1:10 MAPC:Treg on day 0 of each round immediately prior to CD3/28 stimulation. To prepare MAPC cells, the cells were thawed and washed once (500 × g/ 5 mins), then resuspended in complete Treg media. MAPC cells adhered to flat bottom plates and flasks and were absent from final preparations of MulTreg lines as confirmed by microscopy and flow cytometry using anti-CD105 staining on ungated events (data not shown). Following 30 days plus 48h withdrawal, yields were determined by the average of five counts from a single cell suspension and fresh Treg/MulTreg lines were analyzed by flow cytometry or cryopreserved. Sorted Teff cells were cryopreserved without expansion at day 0.

### Flow cytometry

Dead cells were excluded with Fixable Live/dead blue (UV450 Molecular probes, Invitrogen, Life technologies) or 7AAD (BD). Bespoke overlapping Treg panels were constructed using the fluorochrome-labeled antibodies listed in the Supplementary Methods (BD, Biolgend). Intracellular staining was performed using the FoxP3 staining kit (eBiosciences) according to the manufacturers’ instructions. Flow cytometry acquisition was conducted using a BD LSR Fortessa (BD) or BD Celesta (BD), cell sorting completed using the BD FACS ARIA, all equipped with FACS Diva software (v6.0-8.0 (BD Biosciences). Data was analyzed using Flowjo X (Treestar, Ashland, OR).

### Data analysis

Data was analyzed using Prism v8-9 software (Graphpad), and after checking normal distribution of the data, the appropriate statistical test was used for parametric or non-parametric calculations (indicated in the figure legends). Data are shown as mean ± SEM and p values of ≤ 0.05 were considered statistically significant.

### Suppression assays

Cryopreserved Tregs, MulTregs and autologous Teff cells were thawed, labeled with 1μM DDAO (Treg/MulTreg) or Cell trace violet (CTV, Teff) and co-cultured in 96 well plates at the ratios indicated in the figure legends in the presence or absence of CD3/28 (1:50 bead to cell) for 5 days. Proliferation was measured by CTV dye-dilution within viable DDAO^−^ CTV^+^ lymphocytes. Suppression was calculated relative to the proliferation in the absence of Treg or MulTreg cells. Alternatively, cryopreserved Tregs, MulTregs and allogeneic PBMCs were thawed, labeled with 10μM CPDe450 (Treg/MulTreg) or carboxyfluorescein succinimidyl ester (CFSE) (PBMC) and co-cultured in 96 well plates at the ratios indicated in the figure legends in the presence or absence of CD3/28 (Life Technologies) (1:10 bead to cell) for 5 days. Proliferation was measured by CFSE dye-dilution within viable CPDe450^−^ CFSE^+^ lymphocytes. Suppression was calculated relative to the proliferation in the absence of Treg or MulTreg cells. In another experiment, CTV-labeled PBMC were stimulated with 100ng Influenza Hemagglutinin (Flu-HA, Revaxis) for 6 days in the presence or absence of autologous Treg or MulTreg cells and proliferation measured by dye-dilution within CD4^+^ CD3^+^ viable lymphocytes. Cytokine abundance in the supernatant of cell cultures was measured by the LEGENDplex^TM^ multi-analyte flow assay kit, human Th cytokine mix & match subpanel (Biolegend).

### RNA-seq analysis

4 matched pairs of Treg and MulTreg were expanded and RNA extracted (RNeasy RNA extraction kit, Qiagen). cDNA was then produced (SMART cDNA synthesis kit, Clontech) and DNA libraries produced using the Nextera XT kit (Illumina). Library quality was next determined by Bioanalyzer and sequencing performed by MiSeq using v3 chemistry and 150 cycle paired end reads. Sequencing reads generated from the Illumina platform were assessed for quality and trimmed for adapter sequences using TrimGalore! v0.4.2 (Babraham Bioinformatics), a wrapper script for FastQC and cutadapt. Reads that passed quality control were then aligned to the human reference genome (GRCh37) using the STAR aligner v2.5.1. The alignment for the sequences were guided using the GENCODE annotation for hg19. The aligned reads were analyzed for differential expression using Cufflinks v2.2.1, a RNASeq analysis package which reports the fragments per kilobase of exon per million fragments mapped (FPKM) for each gene. Differential analysis report was generated using Cuffdiff. Differential genes were identified using a significance cutoff of q-value < 0.05. The genes were then subjected to gene set enrichment analysis (GenePattern, Broad Institute) to determine any relevant processes that may be differentially over represented for the conditions tested. Additional custom gene set enrichment analysis and visualization were performed in R using the DOSE, enrichplot, and fgsea packages.

#### STAR Aligner

Dobin A, Davis CA, Schlesinger F, Drenkow J, Zaleski C, Jha S, Batut P, Chaisson M, Gingeras TR. STAR: ultrafast universal RNA-seq aligner. Bioinformatics. 2013 Jan 1;29(1):15-21. doi: 10.1093/bioinformatics/bts635. Epub 2012 Oct 25. PubMed PMID: 23104886; PubMed Central PMCID: PMC3530905.

#### Cufflinks

Trapnell C, Williams BA, Pertea G, Mortazavi A, Kwan G, van Baren MJ, Salzberg SL, Wold BJ, Pachter L. Transcript assembly and quantification by RNA-Seq reveals unannotated transcripts and isoform switching during cell differentiation. NatBiotechnol. 2010 May;28(5):511-5. doi: 10.1038/nbt.1621. Epub 2010 May 2. PubMed PMID: 20436464; PubMed Central PMCID: PMC3146043.

### Xenogeneic GVHD model

*Mice:* Immunodeficient NOD.Cg-Prkdcscid Il2rgtm1Wjl/SzJ (NSG) mice were purchased from Charles River Laboratories and maintained in a specific pathogen-free facility (Biological Services Unit, New Hunt’s House, King’s College London). All procedures were performed under sterile conditions in accordance with institutional guidelines and the Home Office Animals Scientific Procedures Act (1986) (Home Office license number: PPL 70/7302). *Xeno-graft vs host disease model:* 6-7-week-old NSG mice were injected intravenously with 1×10^7^ 3^rd^ party PBMCs ± Tregs in a 1:1 ratio. Control mice received saline alone. Following cell transfer, mice were monitored every 2-3 days for signs of xeno-GvHD. Parameters measured included weight loss, hunching, reduced mobility, ruffled hair and orbital inflammation which were graded on a scale of 0-2 [51]. Xeno-GvHD development/progression was scored in a blinded manner by two investigators and mice were sacrificed when pre-defined end-points were reached including >15% weight loss and/or a summed clinical severity score of ≥7. Human CD45+ cell engraftment was measured by flow cytometry in peripheral blood obtained from tail bleeds every two weeks and in the spleen following euthanasia. Mice were considered successfully engrafted and included in analyses when human CD45+ cells constituted >80% of total splenic lymphocytes

### TSDR analysis

Genomic DNA was isolated from d30 post-expansion Treg/MulTreg cells and d0 pre-expansion, sorted Teff cells using the Purelink^TM^ Genomic DNA mini kit (Invitrogen, Life Technologies). Subsequently, 1μg of gDNA was sent to EpigenDx for bisulfate conversion and pyrosequencing to assess the methylation status of the *FOXP3* Treg-specific demethylated region (TSDR). The average methylation percentage of 9 CpG islands was calculated.

## Supplementary material

**Supplementary Table1:** Patient characteristics

|  | **Disease** | **Company/**<br>**Source** | **Gender** | **Age** |
| --- | --- | --- | --- | --- |
| 1 | Crohn's Disease | Precision for Medicine | Male | 25 |
| 2 | Crohn's Disease | Precision for Medicine | Female | 52 |
| 3 | Crohn's Disease | Stem Cell Technologies | Female | 39 |
| 4 | Crohn's Disease | Stem Cell Technologies | Male | 50 |
| 5 | Type1 Diabetes | GSTT NHS foundation trust | Male | 43 |
| 6 | Type1 Diabetes | GSTT NHS foundation trust | Female | 25 |

**Supplementary Table 2:** Ab panel Treg sort

| **Ag** | **fluorochrome** | **Company** |
| --- | --- | --- |
| CD127 clone A019D5 | AF488 | Biolegend |
| CD25 clone 2A3<br>clone MA251 | PE<br>PE | BD Biosciences<br>BD Biosciences |
| CD14 clone 63D3 | PE-Cy7 | Biolegend |
| CD4 clone RPA-T4 | APC | Biolegend |

**Supplementary Table 3:**
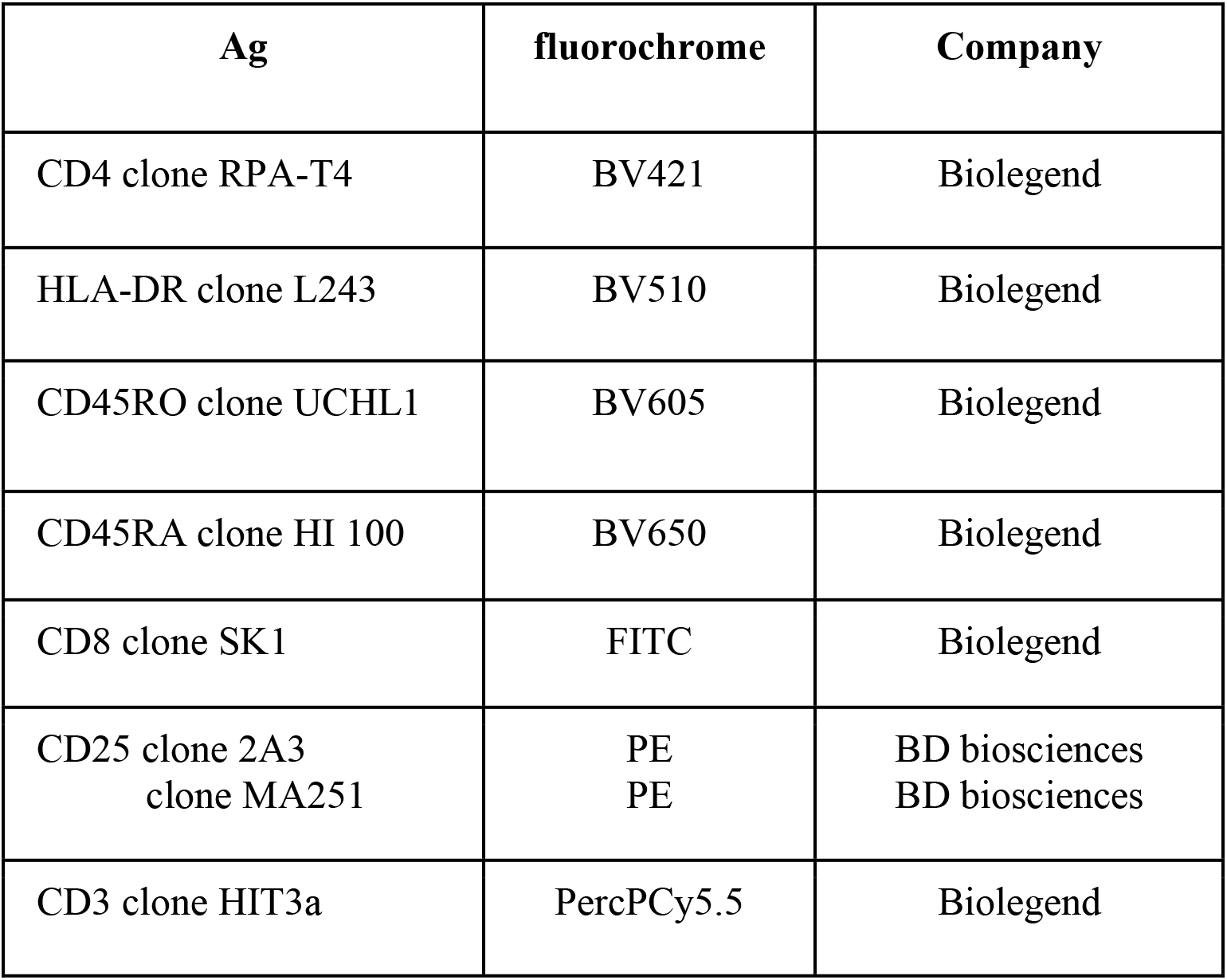
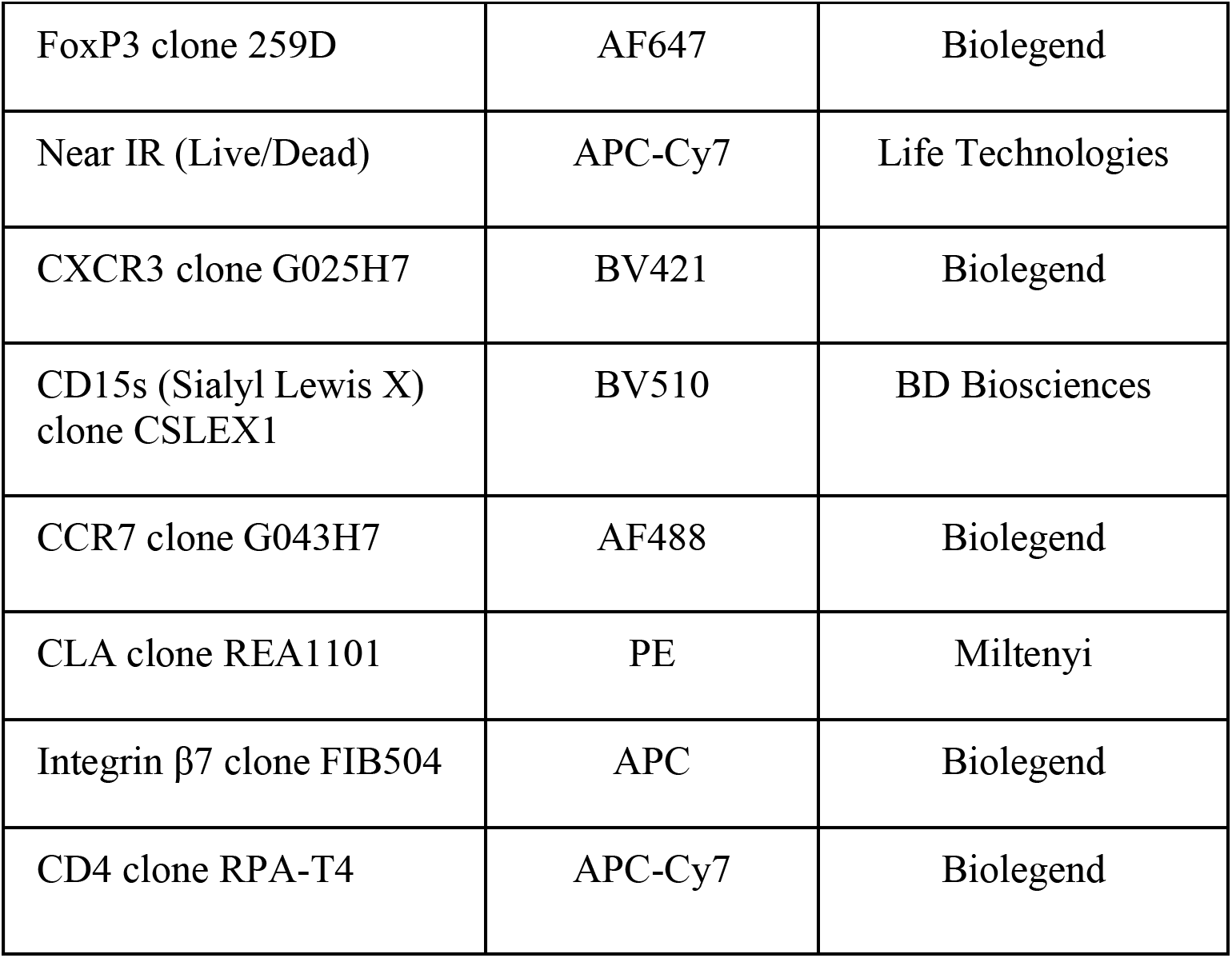
Ab panel Treg characterization

| **Ag** | **fluorochrome** | **Company** |
| --- | --- | --- |
| CD4 clone RPA-T4 | BV421 | Biolegend |
| HLA-DR clone L243 | BV510 | Biolegend |
| CD45RO clone UCHL1 | BV605 | Biolegend |
| CD45RA clone HI 100 | BV650 | Biolegend |
| CD8 clone SK1 | FITC | Biolegend |
| CD25 clone 2A3<br>clone MA251 | PE<br>PE | BD biosciences<br>BD biosciences |
| CD3 clone HIT3a | PercPCy5.5 | Biolegend |
| FoxP3 clone 259D | AF647 | Biolegend |
| Near IR (Live/Dead) | APC-Cy7 | Life Technologies |
| CXCR3 clone G025H7 | BV421 | Biolegend |
| CD15s (Sialyl Lewis X)<br>clone CSLEX1 | BV510 | BD Biosciences |
| CCR7 clone G043H7 | AF488 | Biolegend |
| CLA clone REA1101 | PE | Miltenyi |
| Integrin β7 clone FIB504 | APC | Biolegend |
| CD4 clone RPA-T4 | APC-Cy7 | Biolegend |

## Acknowledgements

We thank Dr. E. Ricky Chan for his assistance with the bioinformatic analysis and Liesbeth Vanenpoel and Ellen Van Houtven for technical assistance.

This research was funded/supported by the National Institute for Health Research (NIHR) Biomedical Research Centre based at Guy’s and St Thomas’ NHS Foundation Trust and King’s College London and/or the NIHR Clinical Research Facility. The views expressed are those of the author(s) and not necessarily those of the NHS, the NIHR or the Department of Health

AET and AV-T are employees of Athersys Inc. and JB and VR are employees of ReGenesys BV a subsidiary of Athersys Inc. AET, AV-T, JB and VR have compensated stock options from Athersys, Inc.

## Author contributions

TT, AT, RD and JR conceived and designed the project. TT, AT, GL and JR supervised the project. JR, CH, VR, JB, ELN, DB, PB designed and executed experiments. All authors interpreted and analyzed data. JR, VR, CH, AVT, AT and TT wrote the manuscript.

---

**Figure S1.**
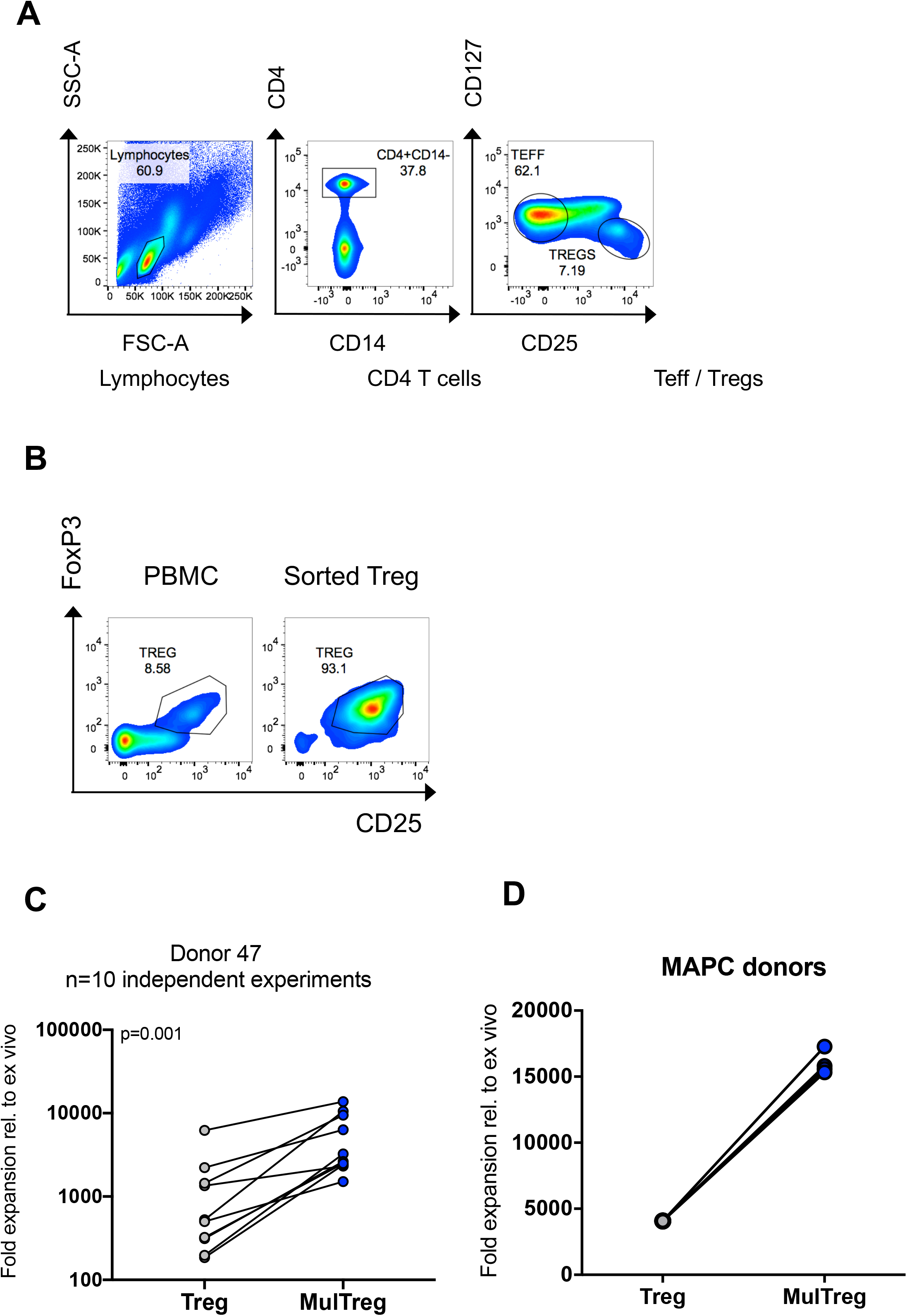
Details of Treg and MulTreg expansion. (A) Identification of Treg and Teff cells sorted analysis and expansion. (B) Example of isolated Treg cells from parent PBMC post sort, prior to expansion. (C)The fold expansion in biological replicates of Treg and MulTreg lines grown from 1 PBMC and 1 MAPC donor (n=10). (D)The fold expansion in Treg and MulTreg lines grown from 1 PBMC donor using 4 different MAPC donors (p=0.065). Stats fromWilcoxon matched-paired signed rank test.

**Figure S2.**
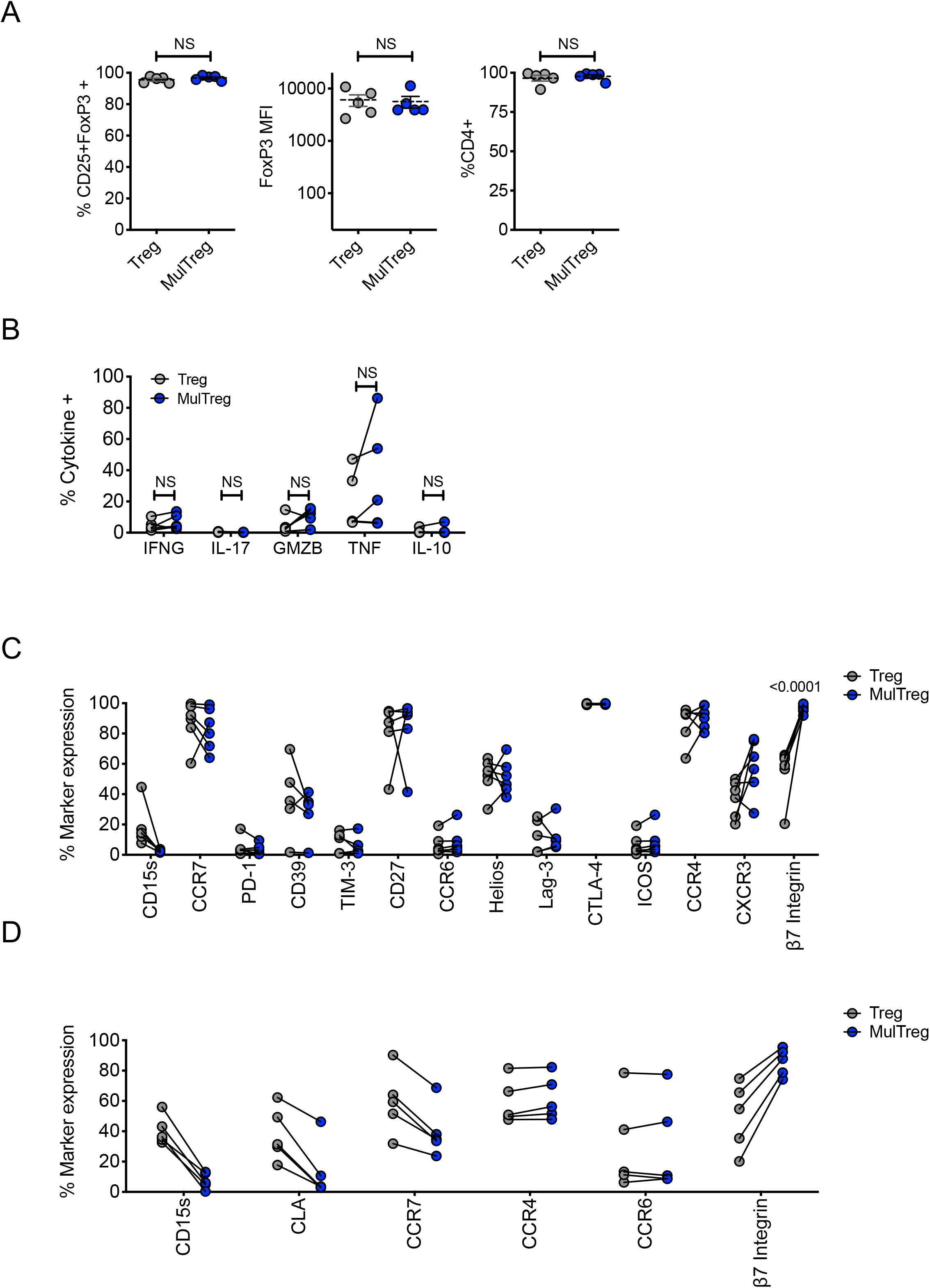
Details of the expanded MulTreg lines. A) Purity of Treg and MulTreg lines in the Belgian cohort. Frequency of CD4+ T cells expressing FoxP3 and CD25 (left), median fluorescence intensity of FoxP3 (centre), frequency of CD4+ T cells (right) in live gated events (n=5). (B) Teff cytokine production in the KCL cohort, n=5 (C) Paired analysis of marker frequencies in the KCL cohort (n=10). (D) Paired analysis of marker frequencies in the Belgian cohort (n=5). All stats from multiple correction adjusted Wilcoxon matched-pairs signed rank test, ns unless shown.

**Figure S3.**
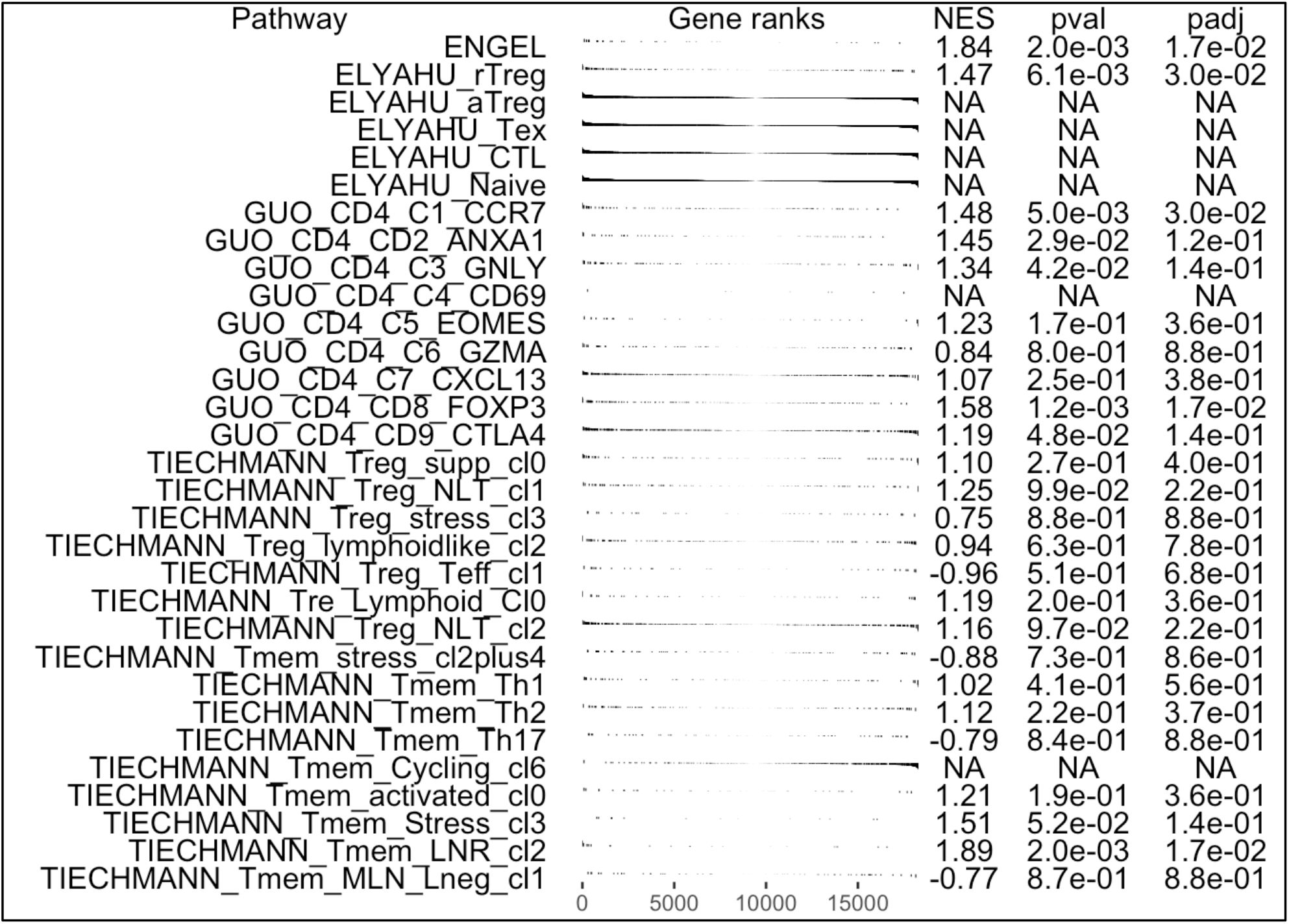
Gene set enrichment analysis (GSEA) of Treg vs MulTreg. All gene sets analysed by GSEA are shown with enrichment plots (Treg vs MulTreg left to right), net enrichment score (NES) with individual and adjusted p values.

**Figure S4.**
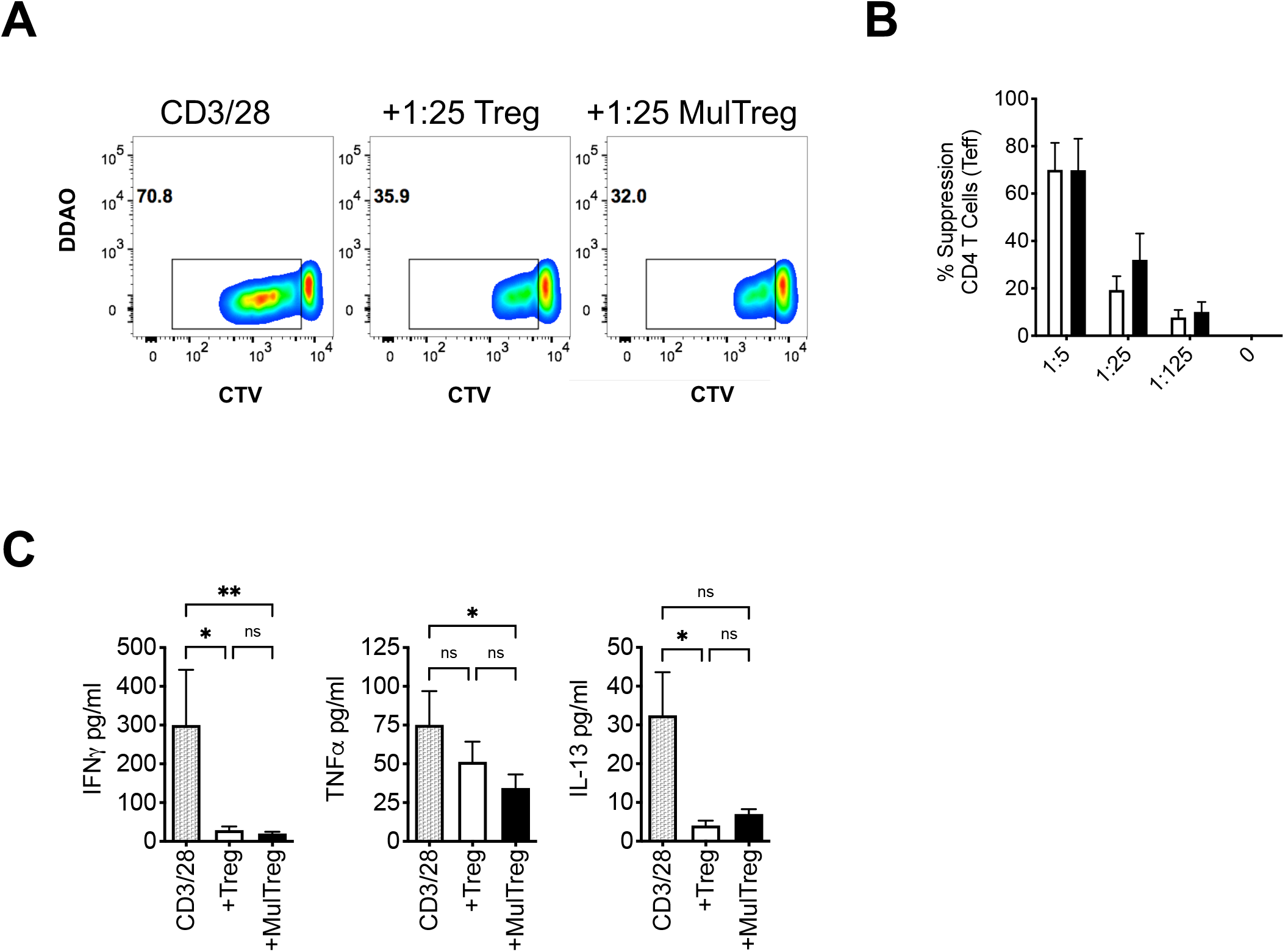
Suppression assay using autologous PBMC stimulated with CD3/CD28 microbeads in the presence or absence of Treg or MulTreg lines. (A). Flow cytometry plots showing proliferation in responder CD4+ T cells from PBMC stimulated with anti-CD3/CD28 for 6 days in the presence or absence of autologous, expanded Treg or MulTreg lines at a ratio of 1:25 Treg:PBMC. (B) Bar graph displaying % suppression of responder CD4+ T cell proliferation at different ratios of Treg:PBMC. (C) Bar graph displaying cytokine levels in tissue culture supernatant from suppression assays in panel B (at ratio of 1:5). Error bars represent the SEM of 6 donors. *p<0.05, **p<0.01

